# The concerted action of E2-2 and HEB is critical for early lymphoid specification

**DOI:** 10.1101/465765

**Authors:** Thibault Bouderlique, Lucia Peña Perez, Shabnam Kharazi, Miriam Hils, Xiaoze Li, Aleksandra Krstic, Ayla De Paepe, Christian Schachtrup, Charlotte Gustafsson, Dan Holmberg, Kristina Schachtrup, Robert Månsson

**Affiliations:** Center for Hematology and Regenerative Medicine, Department of Laboratory Medicine, Karolinska Institutet, Stockholm, Sweden; Center for Chronic Immunodeficiency (CCI), Medical Center, University of Freiburg, Faculty of Medicine & Faculty of Biology, Freiburg, Germany; Institute of Anatomy and Cell Biology, University of Freiburg, Faculty of Medicine, Freiburg, Germany; Lund University Diabetes Center, Lund University, Malmö, Sweden; Hematology Center, Karolinska University Hospital, Stockholm, Sweden

## Abstract

The apparition of adaptive immunity in *Gnathostomata* correlates with the expansion of the E-protein family to encompass E2-2, HEB and E2A. Within the family, E2-2 and HEB are more closely evolutionarily related but their concerted action in hematopoiesis remains to be explored. Here we show that the combined disruption of E2-2 and HEB results in failure to express the early lymphoid program in CLPs and a near complete block in B-cell development. In the thymus, ETPs were reduced and T-cell development perturbed, resulting in reduced CD4 T- and increased γδ T-cell numbers. In contrast, HSCs, erythro-myeloid progenitors and innate immune cells were unaffected showing that E2-2 and HEB are dispensable for the ancestral hematopoietic lineages. Taken together, this E-protein dependence suggests that the appearance of the full *Gnathostomata* E-protein repertoire was critical to reinforce the gene regulatory circuits that drove the emergence and expansion of the lineages constituting humoral immunity.

---

Large *Eumetazoan* rely on an efficient system of innate and adaptive immune cells to survive and reach reproductive age ^1-3^. The different cells of the hematopoietic system are all generated from hematopoietic stem cells (HSCs) ^4^. Lymphoid specification is initiated in lymphoid primed multipotent progenitors (LMPPs) that start to express genes associated with adaptive immune cells ^5,6^. LMPPs subsequently give rise to common lymphoid precursors (CLP) ^7^. Within the heterogeneous CLP population, the LY6D^+^ fraction is further specified towards a B-lineage fate ^8,9^ and contains the first B-lineage committed cells that subsequently give rise to mature B-cells ^10,11^. Early lymphoid precursors leave the bone marrow to seed the thymus where they further develop into early T-cell progenitors (ETP) that give rise to mature T-cells ^12^. Similarly, the innate immune cells develop from different progenitors within the myeloid branch, developing from granulocyte-macrophage progenitors (GMPs) ^13^, while natural killer (NK) cells and part of the dendritic cells (DC) develop from the CLP ^14,15^.

The origin of the *Gnathostomata* (jawed vertebrate) hematopoietic system can be traced far back in evolutionary history with phagocytic and cytotoxic innate immune cells being found across the *Bilateria* ^16^ and the erythroid/megakaryocyte lineages appearing in the *Agnatha* ^17^. Similarly, lymphoid-like cells are present in the *Agnatha* ^18^, *Urochordata* ^19^ and *Cephalochordata* ^20^. However, while genes intimately associated with adaptive immunity - including RAG ^21,22^, histocompatibility genes ^23,24^ and immune type receptors ^23,25,26^ - are found in lower *Deuterostomata*, B- and T-cells mediated adaptive immunity emerged only in the *Gnathostomata*. The appearance of new transcription factor (TF) genes drive the apparition of novel cell types ^27^. The appearance of adaptive immunity in the *Gnathostomata* correlates with a dramatic increase in TF genes ^1,28^. As part of this expansion, the full *Gnathostomata* basic helix-loop-helix E-protein family ^29,30^ consisting of E2A (Tcf3), HEB (Tcf12) and E2-2 (Tcf4) emerged.

It has been proposed, that E2A is more closely related to the ancestral E-proteins while E2-2 and HEB are less evolutionarily conserved and display expression patterns more restricted to vertebrate-specific structures ^30,31^. This suggests that E2A should govern ancestral functions while HEB and E2-2 should govern novel functions that emerged concomitantly to the rise of the *Gnathostomata*. In line with this, E2A is the only E-protein reported to impact HSC function and the development of the myeloid- and erythro/megakaryocytic lineages ^32,33^. In contrast, all the E-proteins promote development of B- and T-cells ^9,33-43^. The potential role of E2-2 in stem- and progenitor cells remains largely unexplored.

Here we confirm that E2-2 and HEB are evolutionary related and we found that their coordinated action is critical for the development of early lymphoid progenitors. Mice lacking both E2-2 and HEB display an almost complete block in B-cell development at the level of the CLP and the few generated immature B-cells preferentially develop into marginal zone (MZ) B-cells. Similarly, we found T-cell development to be perturbed, resulting in reduced numbers of CD4 T-cells and increased numbers of γδ T-cells. In contrast, HSCs, erythro-myeloid development and the generation of innate immune cells were unperturbed. Together, this suggests that E2-2 and HEB are dispensable for ancestral hematopoietic lineages and that the appearance of the full *Gnathostomata* E-protein repertoire promoted the apparition of humoral immunity.

## Results

### E2-2 and HEB are evolutionarily related

To investigate the evolutionary history of the E-proteins in *Bilateria* in light of recent sequencing data, we analyzed similarities between cDNA and amino acid sequences of the E-proteins across *Animalia* (Fig. 1A). The E-proteins of *Protostomata* and non-gnathostome *Deuterostomata* clustered together (Fig. 1B) and displayed relatively high sequence divergence (Fig. 1C) on the cDNA level. The *Gnathostomata* E-proteins, in contrast, formed a separate clade (Fig. 1B) and displayed comparably higher similarity (Fig. 1C). Within the *Gnathostomata* clade, E2-2 and HEB formed a separate branch from that of E2A (Fig. 1C). Similar results were obtained from the analysis of the amino acid sequences (Fig. S1). While only providing weak support for E2A being more closely related to the ancestral E-protein, it confirms a closer evolutionary relation between E2-2 and HEB ^30^. It could hence be hypothesized that E2-2 and HEB are functionally related and together support the development of cell lineages specific to the jawed vertebrate hematopoietic system.

**Figure 1.**
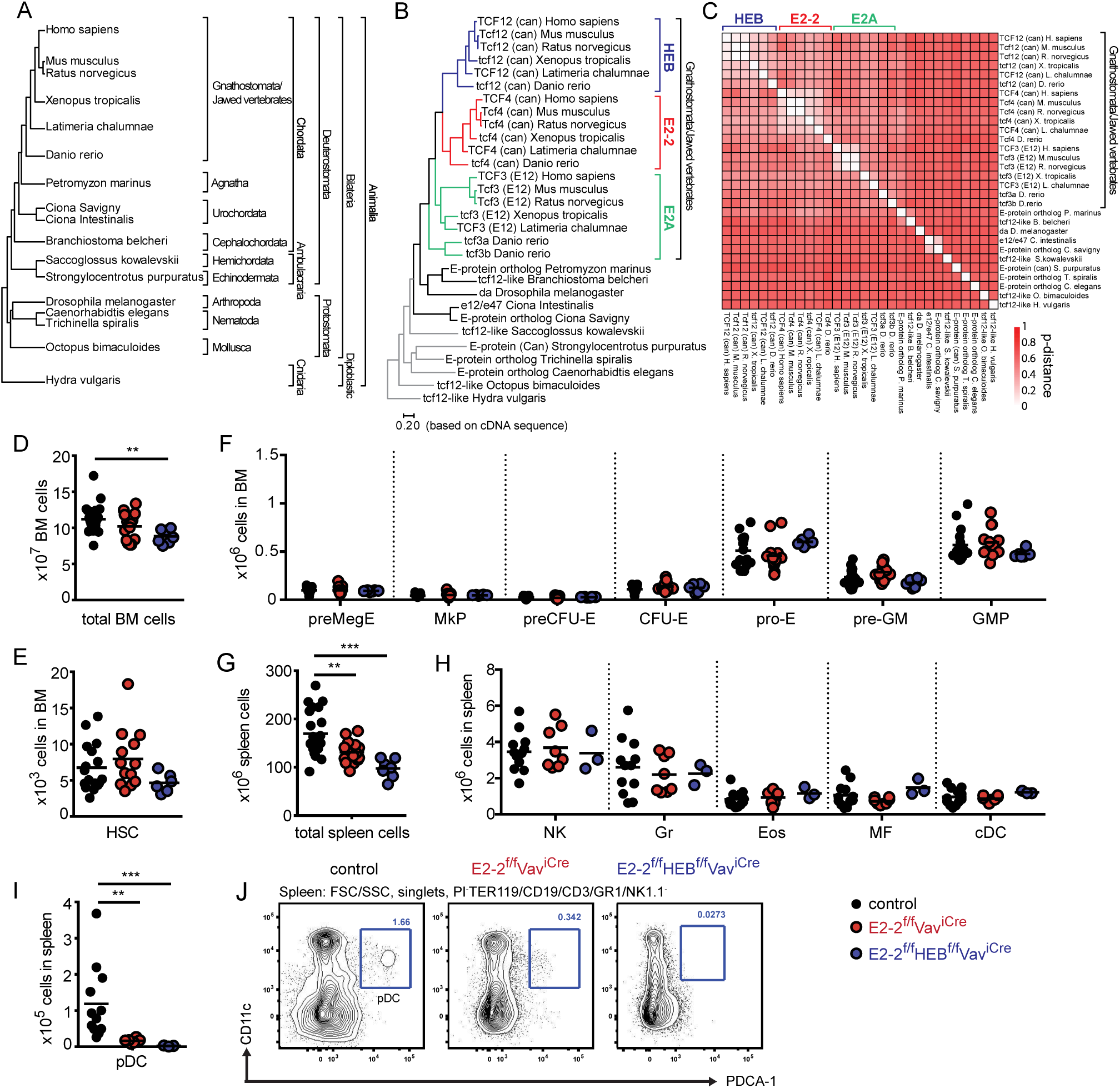
E2-2 and HEB are evolutionarily related and dispensable for ancestral lineages. (A) Schematic overview of evolutionary relationships and classification of animals included in the E-protein phylogenetic analysis. (B) Maximum Likelihood phylogenetic tree of the evolutionary distances between E-protein cDNA sequences in *Bilateria*. *Hydra vulgaris* was used as an out-group for the analysis. Colors indicate the E-protein families in *Gnathostomata*. Grey indicates branches with low support in the bootstrapping analysis (<70% of trees generated maintain the branches). For sequences used in the analysis, see Table S1. (C) Matrix displaying the pairwise p-distances between analyzed cDNA sequences. Absolute number of total cells (D), hematopoietic stem cells (HSCs) (E) and erythro-myeloid progenitors (F) in bone marrow. Absolute number of total cells (G), innate immune cells (H) (granulocytes, Gr; eosinophiles, Eos; macrophages, MF; and conventional dendritic cells, cDC) and plasmacytoid dendritic cells (pDC) (I) in spleen. For gating strategies for cell types in panel EF and H see Fig. S3A-C. (J) Gating strategy for identification of pDCs. Color symbols utilized throughout the figure to indicate the genotype of analyzed mice are shown in the bottom right corner. Significance was calculated using the Mann-Whitney U test with *, ** and *** indicating p-values <0.05, <0.01 and <0.001 respectively.

### Deletion of E2-2 and HEB does not impact HSCs and erythro-myeloid progenitors

To investigate the role of E2-2 and HEB in hematopoiesis, we used the Vav^iCre^ mouse strain to mediate conditional deletion of E2-2 (E2-2^f/f^Vav^iCre^) or E2-2 together with HEB (E2-2^f/f^HEB^f/f^Vav^iCre^) in the hematopoietic system. Cre mediated deletion of the floxed exons was verified in RNAseq data from FACS sorted cells (Fig. S2A-C). We observed a difference in bone marrow (BM) cellularity only between littermate control mice (lacking Vav^iCre^) and E2-2^f/f^HEB^f/f^Vav^iCre^mice (Fig. 1D). However, no differences were observed in the number of HSCs (Fig. 2E, S3A), megakaryocyte, erythroid and myeloid progenitors (Fig. 2F, S3B). While being expressed in the HSC and erythro-myeloid progenitors (Fig. S2D), E2-2 and HEB are hence dispensable for the maintenance and generation of these cell types in steady state ^33^. This leaves E2A as the sole E-protein needed for HSCs and erythro-myeloid development ^32^.

**Figure 2.**
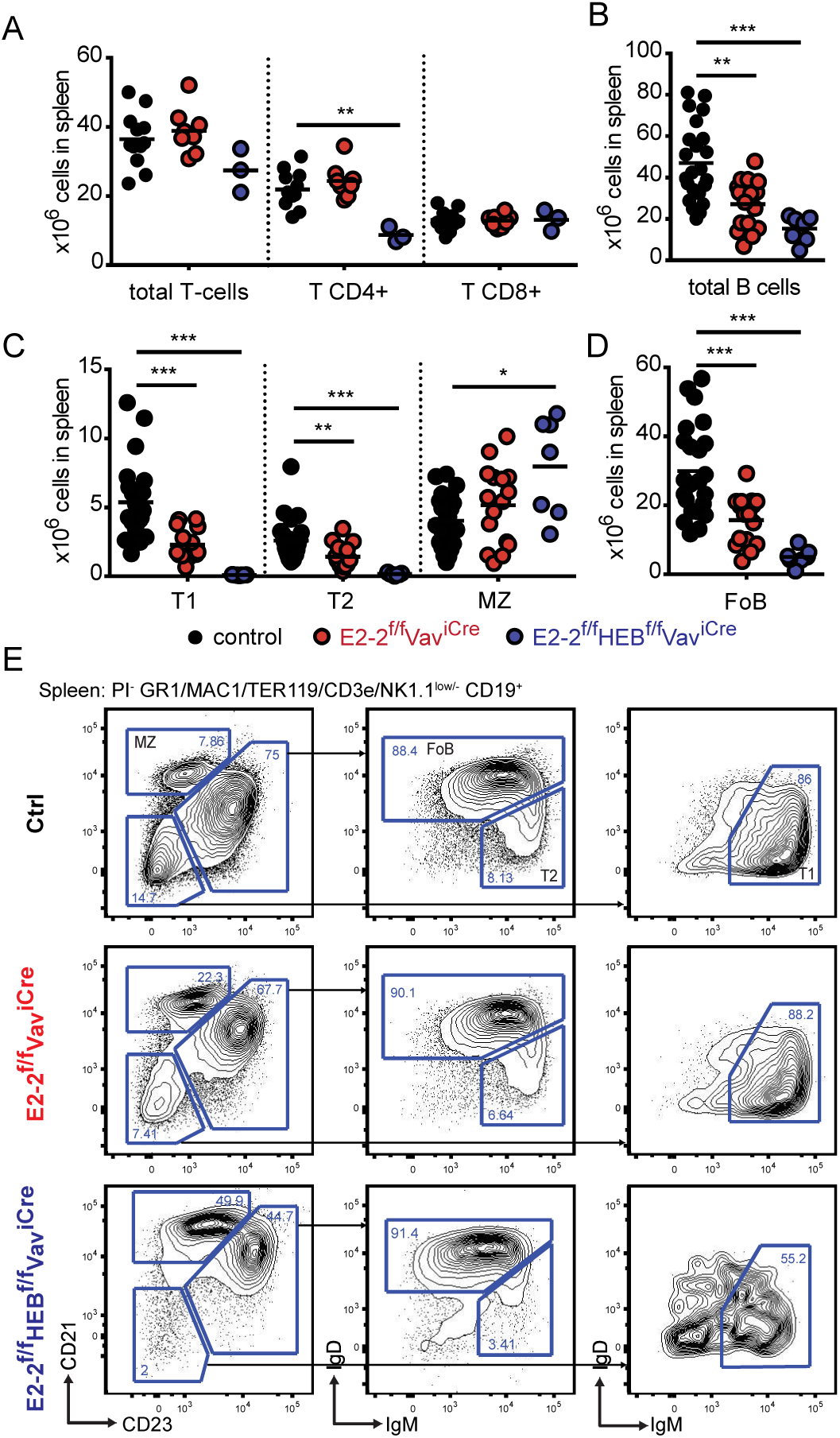
E2-2 and HEB are needed for proper B-cell and CD4 T-cell generation. (A) Absolute number of T-cells, CD4 T-cells and CD8 T-cells in spleen. For T-cell gating strategy see Fig. S3C. Absolute number of B-cells (B) and B-cell subsets (C-D) (including: transitional 1-, T1; transitional 2-; marginal zone-, MZ; and follicular B-cells, FoB) in spleen. (E) Gating strategy for identification of B-cell subsets. Color symbols utilized throughout the figure to indicate the genotype of analyzed mice are shown below panels C-D. Significance was calculated using the Mann-Whitney U test with *, ** and *** indicating p-values <0.05, <0.01 and <0.001 respectively.

### Generation of innate immune cells except pDCs are unaffected by E2-2 and HEB deletion

While the total number of spleen cells was reduced in E2-2^f/f^HEB^f/f^Vav^iCre^mice (Fig. 1G), no effect was observed on the number of mature myeloid cells (Fig. 1H and S3C). This further support that E2-2 and HEB are dispensable for myelopoiesis. Similarly, natural killer cell numbers were unaffected (Fig. 1H and S3C). In contrast, plasmacytoid dendritic cells (pDC) were reduced by 86% by the deletion of E2-2 and by 98% with the additional removal of HEB (Fig. 1I-J) indicating a previously unrecognized dependence on HEB for pDC development ^44^.

### E2-2 and HEB are needed for proper generation of B-cells and CD4 T-cells

As the total number of spleen cells was reduced (Fig. 1G), we next investigated the effect of deletion of E2-2 and HEB on the adaptive immune cells. The number of T-cells in the spleen was not impacted by the depletion of E2-2, but E2-2^f/f^HEB^f/f^Vav-iCre mice displayed a 60% reduction in CD4 T-cells (Fig. 2A). The total number of B-cells in spleen was reduced by 42% by the deletion of E2-2 and by 68% when HEB was additionally depleted (Fig. 2B). This reduction reflected a sharp drop in transitional- and follicular B-cells (FoB) while marginal zone (MZ) B-cell numbers remained unaffected in E2-2^f/f^Vav-iCre and increased in E2-2^f/f^HEB^f/f^Vav-iCre mice (Fig. 2C-E). This suggests that the removal of E2-2 and HEB heavily promoted the generation of MZ from transitional B-cells at the expense of FoB ^45,46^. Taken together, these data show that cell lineages with ancestral functions (erythro-myeloid and cytotoxic cells)^16,17,47-50^ are independent of E2-2 and HEB. In contrast, cell lineages central to adaptive (humoral) immunity (B- and CD4 T-cells) or at the interphase between innate and adaptive immunity (pDC) are dependent on E2-2 and HEB.

### The development of CD4 and γδ T-cells is perturbed by the loss of E2-2 and HEB

With E2-2^f/f^HEB^f/f^Vav^iCre^ mice displaying decreased CD4 T-cells in the periphery (Fig. 2A), we next investigated T-cell development in the thymus (Fig. 3A). Depletion of E2-2 had no effect on thymic cellularity (Fig. 3B). However, E2-2^f/f^Vav^iCre^ mice displayed a visible decrease in ETPs and DN2 followed by decreases in DN3E and IS(8)P (Fig. 3D). In line with previous studies of HEB knock-out animals ^34^, the additional deletion of HEB perturbed T-cell development (Fig. 3A) with decreased total cellularity in thymus (Fig. 3B); significant decreases in ETP and DN2 (Fig. 3C); increases in DN3E, DN3L and IS(8)P (Fig. 3D); but markedly reduced DPs (Fig. 3D). In spite of the marked reduction in DPs, CD8 T-cells were present in normal numbers (Fig. 3E) while CD4 T-cells were severely reduced (>85%). Additionally, E2-2^f/f^HEB^f/f^Vav^iCre^ animals displayed markedly increased (6-fold) γδ T-cells numbers (Fig. 3E) ^34,51^.

**Figure 3.**
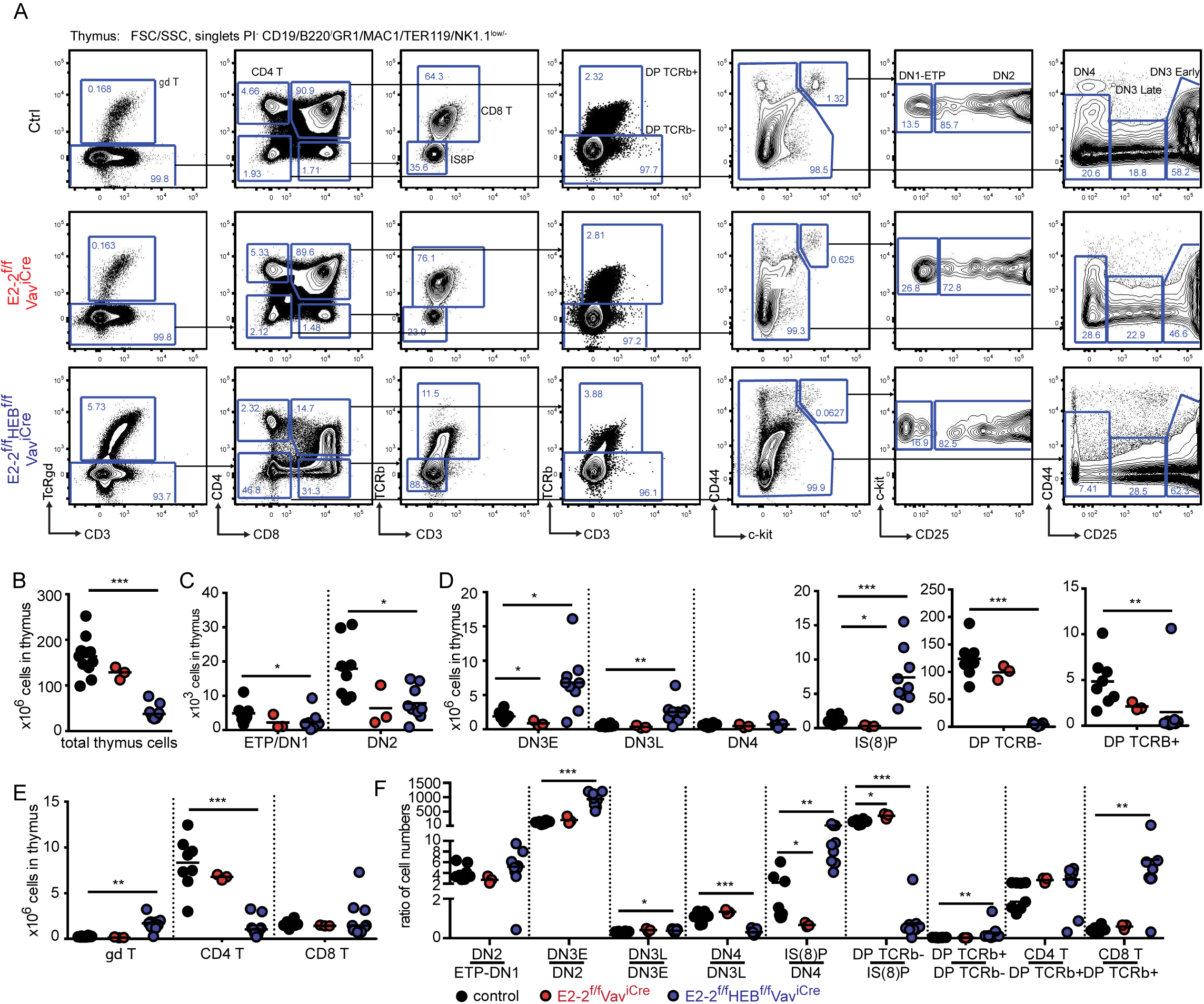
Maturation of CD4 and γδ T-cells is perturbed by the loss of E2-2 and HEB. (A) Gating strategy for identification of T-cell developmental stages. Absolute numbers of: total cells (B); ETP/DN1 and DN2 (C); DN3E, DN3L, DN4, IS(8)P, DP TCRβ^−^ and DP TCRβ^+^ (D); and γδ T-, CD4 T- and CD8 T-cells (E) in thymus. (F) Cell number ratios for each consecutive stage of development as a fraction of the prior stage. Each dot represents the ratio for one individual animal. Color symbols utilized throughout the figure to indicate the genotype of analyzed mice are shown at the bottom of the figure. Significance was calculated using the Mann-Whitney U test with *, ** and *** indicating p-values <0.05, <0.01 and <0.001 respectively.

Looking closer at the progressive generation of cells from the prior stage in each developmental transition in E2-2^f/f^HEB^f/f^Vav^iCre^ mice, the altered cell numbers observed were mirrored by increased generation of DN3E, DN3L and IS(8)P before a sharp drop in the generation of DP TCRβ^−^cells (Fig. 3F). While few in numbers, DP TCRβ^+^ were generated in increased numbers with CD4 and CD8 T-cells subsequently being generated at a normal and increased ratio respectively (Fig. 3F). Taken together these results demonstrate that E2-2 and HEB are critical for thymopoiesis and the normal generation of γδ and CD4 T-cells.

### E2-2 and HEB are critical for lymphoid specification and B-cell development

The development of LMPPs and CLPs constitute the first steps of lymphoid specification ^5,7,11^. Depletion of E2-2 alone did not significantly affect the number of LMPPs and Ly6D^−^CLPs (Fig. 4A-C). However, the number of B-cell specified LY6D^+^CLPs ^8,11^ was reduced by half (Fig. 4C). Correspondingly, total B-cells were reduced by 43% (Fig. 4D) with each stage in B-cell development displaying a 30-50% reduction (Fig. 4E-F). The additional loss of HEB, in contrast, lead to a 70% decrease in the number of LMPPs and LY6D^−^CLP (Fig. 4A-C). In addition, the E2-2^f/f^HEB^f/f^Vav^iCre^ mice strikingly displayed a near complete loss of LY6D^+^CLP (>98% reduction) (Fig. 4A-B). This phenotype is similar to what has previously been reported for E2A knock-out ^33^. Accordingly, B-cell numbers were severely reduced (Fig. 4D) with a >99% reduction in cell-numbers of maturing B-cells (Fig. 4E-F) and a 94% reduction in mature B-cells (Fig. 4E-F) in BM.

**Figure 4.**
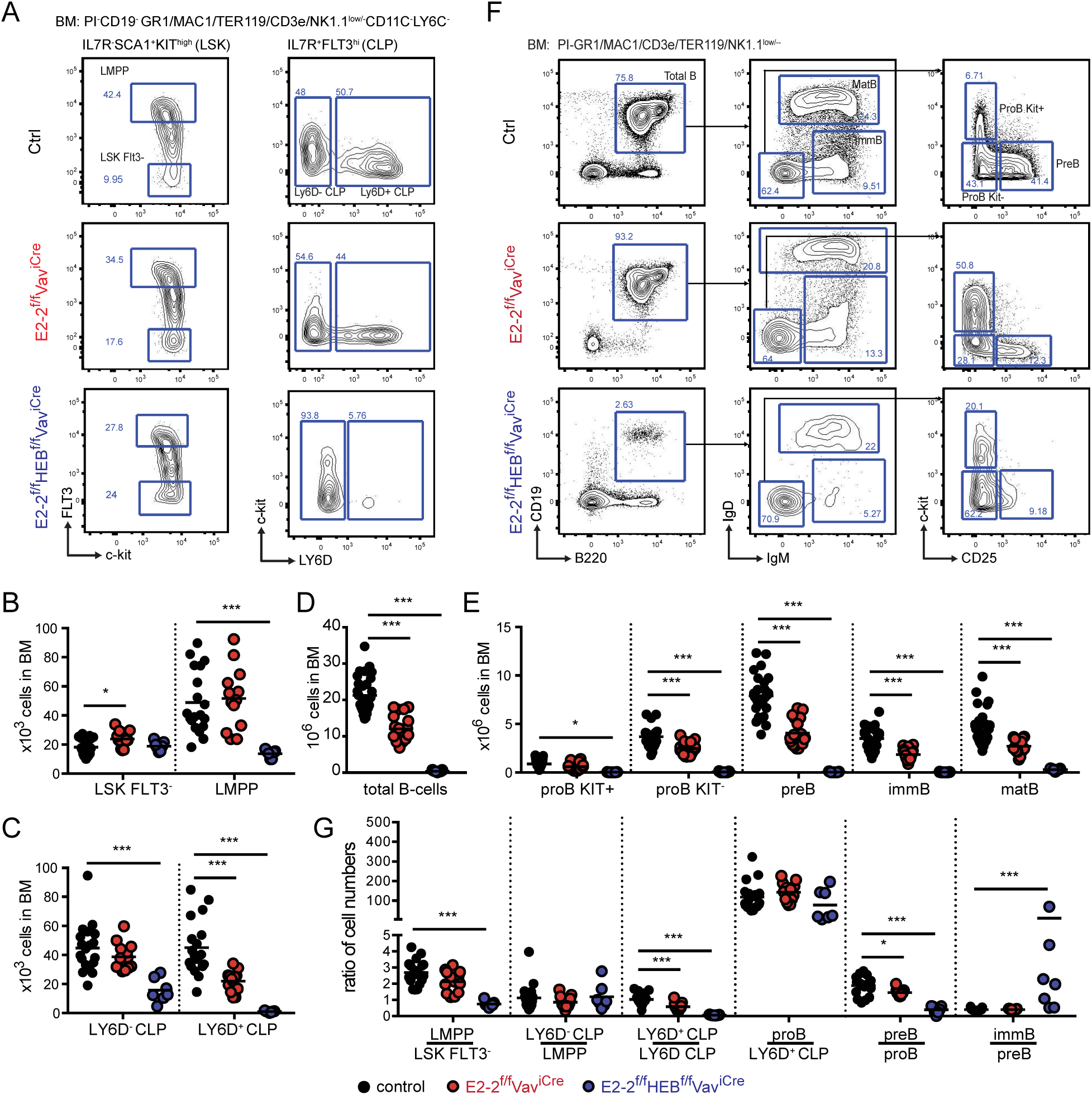
The combined activity of E2-2 and HEB are critical for the generation of lymphoid progenitors and B-lineage cells. (A) Gating strategy for identification of stem- and lymphoid progenitors. (B-C) Absolute numbers of stem- and lymphoid progenitors indicated cell types in BM. Absolute numbers of total B-cells (D) and each B-cell developmental stage (E) in BM. (F) Gating strategy for identification of total B-cells and B-cell developmental stages. (G) Cell number ratios for each consecutive stage of development as a fraction of the prior stage. Each dot represents the ratio in an individual animal. Color symbols utilized throughout the figure to indicate the genotype of analyzed mice are shown below panel G. Significance was calculated using the Mann-Whitney U test with *, ** and *** indicating p-values <0.05, <0.01 and <0.001 respectively.

To analyze more closely the impact of E2-2 and HEB depletion on developmental transitions, we plotted cell number ratios at each consecutive stage of development as a fraction of the prior stage (Fig. 4G). E2-2 deficient mice only displayed a modest decrease in the generation of LY6D^+^CLP from LY6D^−^CLP while the consecutive generation of CD19^+^ developmental stages remained unaffected (Fig. 4G). This indicates that E2-2, similarly to what has been reported for HEB deficient animals ^33^, is important for the LY6D^−^ to LY6D^+^ CLP transition but largely dispensable for BM B-cell maturation. This is in line with the expression data showing that early lymphoid progenitors (LMPPs and CLPs) expressed similar levels of E2-2, HEB and E2A while HEB and E2A were the predominantly expressed E-proteins in B-lineage cells (Fig. S2D). The combined deletion of E2-2 and HEB severely impacted the generation of LMPPs, LY6D^+^CLP and pre-B while, at the same time, seemingly increasing the generation of immature B-cells (Fig. 4G). Interestingly, the generation of proB cells from the few remaining LY6D^+^CLP was unaffected (Fig. 4G), indicating that B-lineage commitment at this stage is unperturbed by the lack of E2-2 and HEB. Together, this demonstrates that the collaboration of E2-2 and HEB is critical for the generation of early lymphoid progenitors and the development of B-cells in the BM.

### Expression of the early lymphoid program is disrupted by removal of E2-2 and HEB

To better understand the mechanisms behind the impaired generation of the B-cell specified LY6D^+^CLPs and to understand how it relates to the similar phenotype observed in mice lacking E2A ^33^, we characterized the transcriptional profiles of LY6D^−^CLPs remaining in E2-2^f/f^Vav^iCre^, E2-2^f/f^HEB^f/f^Vav^iCre^ and E2A^f/f^Vav^iCre^ mice using RNAseq (see Table S2 for sample information). Principal component analysis (PCA) of full expression profiles showed that LY6D^−^CLP cells from the different genotypes form a distinct cluster and hence represent the same population of cells regardless of disrupted E-protein gene(s) (Fig. 5A). A total of 150 genes displayed significant (corrected p-value <0.01) expression changes in LY6D^−^CLPs in one or more of the analyzed strains (Fig. 5B). As expected, B-lineage related genes (including Ebf1, Blnk, Blk, Ets1, Dntt, Notch1, Rag1 and Rag2) were severely affected (Fig. 5B) and overall the gene set was functionally associated with lymphocyte differentiation and signaling (Fig. S4). While the majority of genes did not show highly significant changes in all genotypes (Fig. 5C-D), the overall pattern of the expression changes was highly similar in E2-2^ff^HEB^ff^Vav^iCre^ and E2A^ff^Vav^iCre^ mice (Fig. 5B and D-E). The modest expression changes observed in the E2-2^ff^Vav^iCre^ mice were often concordant to those observed in E2A^ff^Vav^iCre^ mice and E2-2^ff^HEB^ff^Vav^iCre^ mice (Fig. 5B and D-E). Taken together, this shows that E2-2, HEB and E2A largely reinforce the same gene network in early lymphoid progenitors while having variable impact on the expression of individual genes.

**Figure 5.**
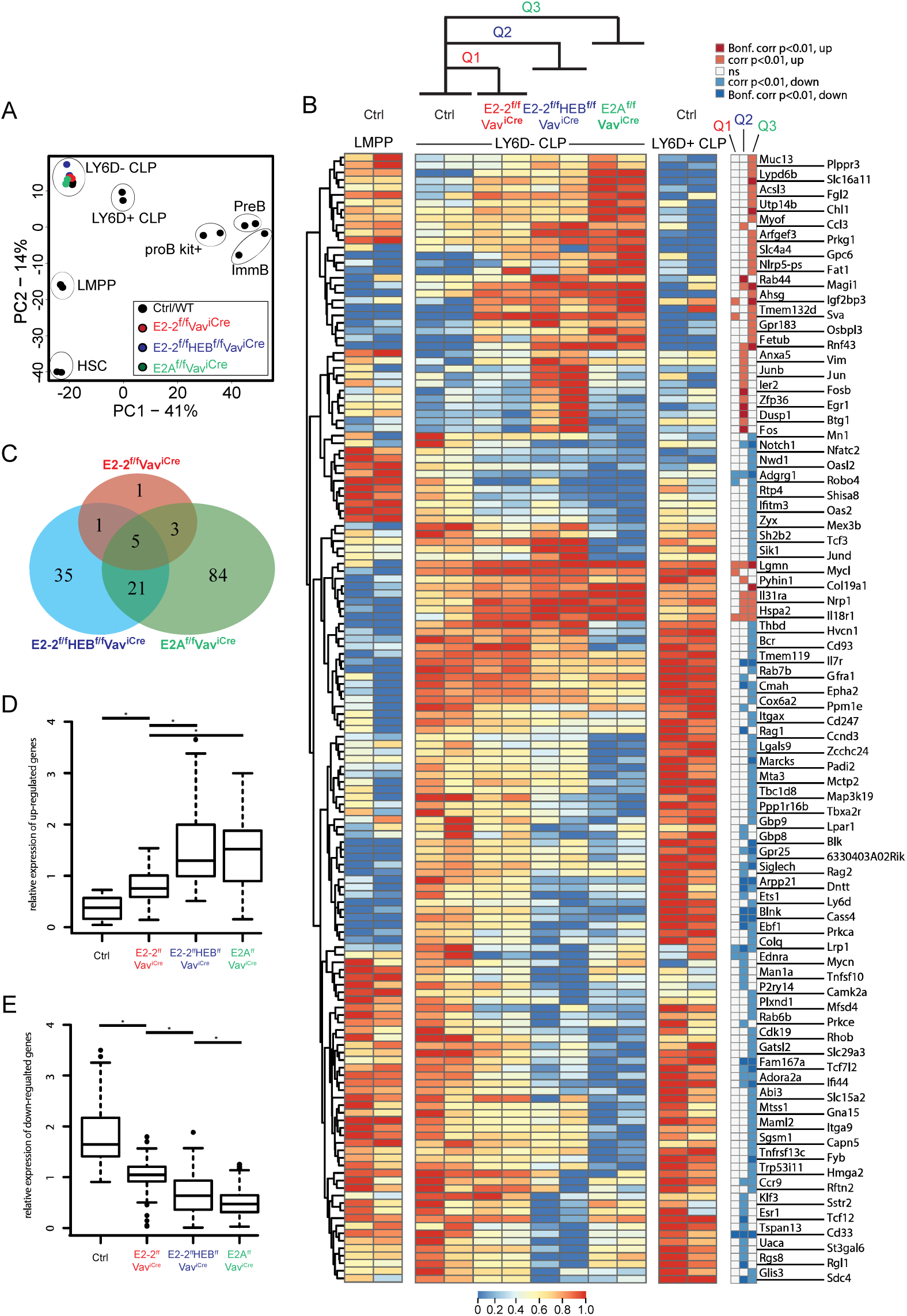
Loss of E-protein activity disrupts the early lymphoid transcriptional program. (A) Principal component analysis of RNAseq data. (B) Hierarchically clustered heatmap showing the (row normalized) expression of genes with significant (adjusted p-value <0.01 calculated by EdgeR) changes in LY6D^−^CLPs from E2-2^f/f^Vav^Cre^ (Q1), E2-2^f/f^HEB^f/f^Vav^Cre^ (Q2) or E2A^f/f^Vav^Cre^ (Q3) as compared to control (ctrl). Significance of expression changes in each comparison (Q1-3) is indicated to the right of the heatmap. (C) Venn diagrams showing the overlap between the significant expression changes in the indicated genotypes. Relative expression change (mean expression is set to one for each gene) of up-regulated (D) and down-regulated genes (E) from Q1-Q3. * indicates a p-value <0.002 calculated using the Mann-Whitney U test.

### E-proteins display partly overlapping association with chromatin

To further understand the mechanisms through which E2-2 and HEB control B lymphopoiesis, we analyzed the binding pattern of E2A, HEB and E2-2 in pro B-cells using ChIP-seq (see Table S2 for sample information). We identified 16510 E2A, 2167 HEB and 139 E2-2 high quality peaks (peak score ≥10) (Fig. 6A and S5A). E2-2 is the lowest expressed E-protein in pro B-cells (Fig. S2D) and the relatively low enrichment of E2-2 (most peaks have a peak score <10, Fig. S5A) limits accurate peak calling. Hence, E2-2 binding is likely underestimated. Peaks from the three E-protein ChIPseq experiments displayed highly significant enrichment of E-protein DNA binding motifs (Fig. S5B).

**Figure 6.**
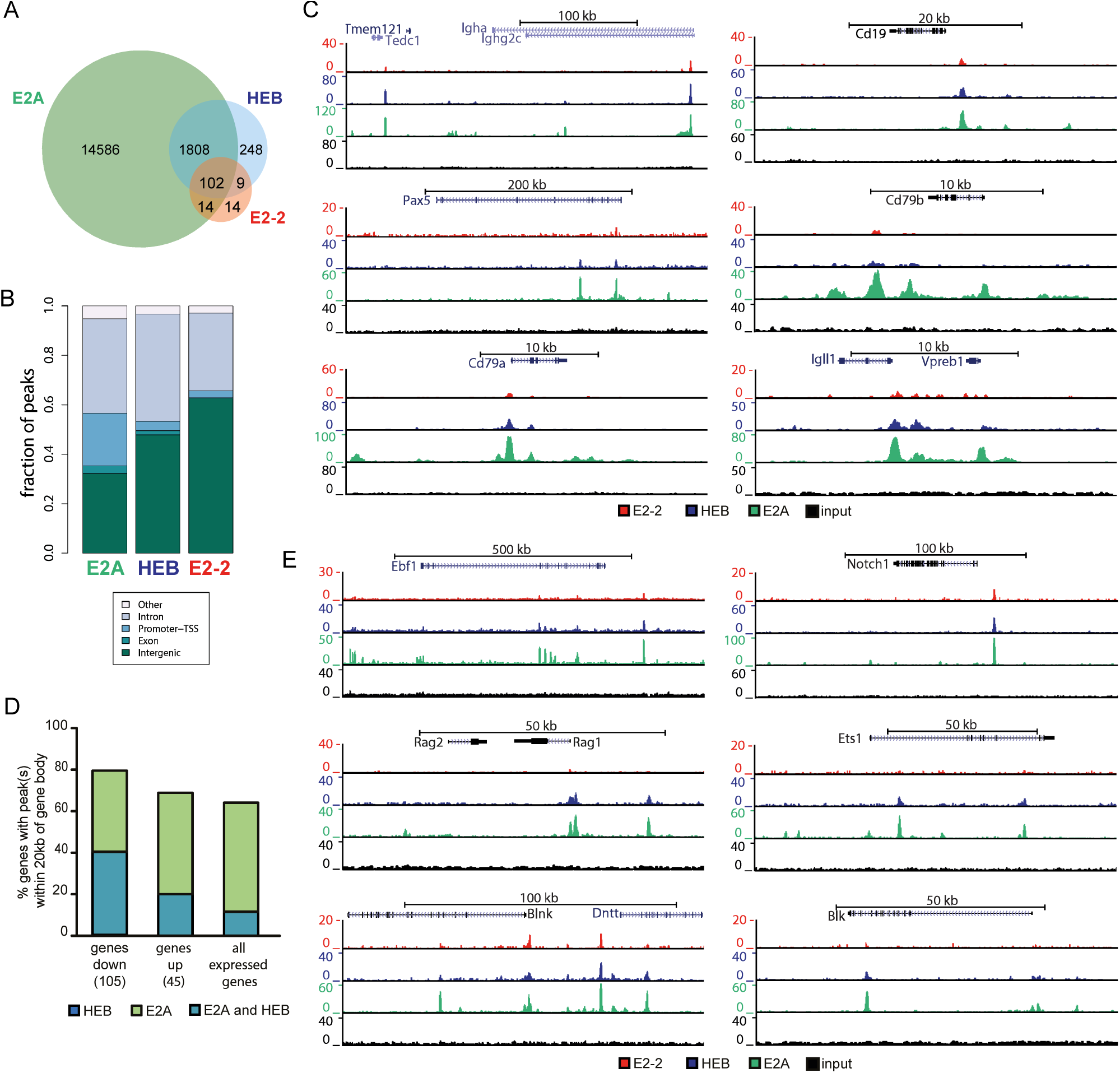
E2-2, HEB and E2A have partly overlapping binding patterns. (A) Number and overlap of identified E2-2, HEB and E2A peaks in proB cells. (B) Genomic localization of identified peaks. (C) Genome browser tracks showing E-protein binding near central B-lineage genes. (D) Percentage of genes in the E-protein dependent early lymphoid program (up/down regulated in Fig. 5B) and all expressed genes in LY6D^−^CLPs that have HEB and E2A peaks within the gene body ±20kb. (E) Genome browser tracks showing E-protein binding near genes that were regulated by E-proteins in LY6D^−^CLPs.

E2A mostly bound regions without identified HEB and E2-2 peaks (Fig. 6A and S5C). However, the majority of E2-2 and HEB peaks overlapped with those of E2A (Fig. 6A and S5C). Most binding sites identified where localized in intronic and intergenic regions (Fig. 6B). In contrast to the other E-proteins, a significant fraction of E2A peaks were also found in promoter regions (Fig. 6B). No difference was observed in the E-protein DNA binding motifs of HEB and/or E2A, with the most enriched motifs containing the same bHLH motif core (CAGCTG) ^52^. The most enriched motif in E2A peaks was present amongst the motifs found in the peaks common to E2A and HEB (Fig. S5D). Genes central to B-lineage, including IgH, CD19, Pax5, CD79a, CD79b, Igll1 and VpreB, were bound by the three E-proteins (Fig. 6C). A similar pattern was found for genes that are part of the E-protein-dependent early lymphoid program in LY6D^−^ CLPs (Fig. 5B), including Ebf1, Rag1, Rag2, Blnk, Dntt, Notch1 and Blk (Fig. 6D-E and S5E). These regions at the center of the lymphoid and B-lineage programs, bound by E2A and HEB, presented a higher enrichment for Ebf and Pax DNA binding motifs compared to regions with only detectable E2A peaks (Fig. S5F).

Taken together this further supports the notion that E2-2 and HEB critically reinforce the action of E2A on gene regulatory circuits critical to early lymphoid specification and B-cell development.

## Discussion

Phylogenetic analysis of the E-proteins suggests that the *Gnathostomata* E-protein family arose through two subsequent duplications ^30,53^. The initial duplication generated two E-protein loci that diverged into the proto-E2-2/HEB and E2A loci. The second duplication subsequently gave rise to the E2-2 and HEB loci as well as two E2A loci out of which one was eventually lost ^53,54^. Being evolutionarily related, E2-2 and HEB could hence support similar functions. The E-proteins play critical roles in all the hematopoietic lineages ^9,32-34,36,38-46,55,56^. However, the function of E2-2 in early hematopoietic development and its concerted action with the evolutionarily related E-protein HEB has not been thoroughly addressed.

The B-cell developmental pathway has been shown to critically rely on E2A and HEB, both TFs needed for the generation of LY6D^+^CLPs ^33^. We similarly found that the loss of E2-2 impaired the generation of LY6D^+^CLPs while earlier progenitors (including LMPPs and LY6D^−^CLPs) were unaffected. Further development of B-cells from LY6D^+^CLPs was unperturbed with the subsequent stages appearing at expected ratios. This indicates that the reduced number of B-cells is primarily caused by the impaired LY6D^−^ to LY6D^+^ transition within the CLP compartment.

Interestingly, the combined deletion of E2-2 and HEB had a direct additive effect resulting in a near complete developmental block at the LY6D^−^CLP stage and dramatically reduced B-cell numbers. Hence, E2-2 and HEB together have a critical and previously unrecognized role in supporting early lymphoid development in the BM. This phenotype is reminiscent of the one observed in E2A knockout animals ^33^. In line with this, the genes of the early lymphoid program were associated with combined E2A and HEB binding. This further supports the notion that all three E-proteins, to a large extent, reinforce the same gene regulatory circuit in CLPs ^33^.

The dramatic reduction of BM B-lymphopoiesis in E2-2^ff^Vav^iCre^ and E2-2^ff^HEB^ff^Vav^iCre^ animals was mirrored in reduced numbers of transitional (T1 and T2) B-cells and reduced FoBs. In contrast MZ B-cell numbers were maintained in E2-2^ff^Vav^iCre^ mice or even expanded in E2-2^ff^HEB^ff^Vav^iCre^mice. This means that MZ B-cells are generated at the expense of FoBs, confirming that the levels of all three E-proteins are critical to maintain a normal MZ to FoB ratio ^45,46^.

E2-2^ff^Vav^iCre^and E2-2^ff^HEB^ff^Vav^iCre^ mice displayed, on average, a 60% reduction in ETPs. Hence, while at a reduced level, E2A alone is sufficient to maintain thymic seeding in the absence of the other E-proteins ^36,42^. T-cell development downstream of the ETP was significantly perturbed only in E2-2^ff^HEB^ff^Vav^iCre^ animals resulting in reduced CD4 T-cells and increased generation of γδ T-cells. This phenotype is similar to the one observed in the HEB KO mice ^34^. Hence, this suggests that E2-2 has limited impact on adult thymopoiesis ^44^.

In contrast to the clear impact of E2-2 and HEB on cells involved in humoral immunity (B- and CD4 T-cells), the loss of these TFs did not affect HSC numbers, erythro-myeloid progenitors nor the production of the major innate immune cells lineages including granulocytes, macrophages and natural killer cells. Similarly the cytotoxic branch of adaptive immunity (CD8 T-cells) was unaffected. Taken together, this suggests that E2-2 and HEB - in contrast to E2A ^32,33^ - are dispensable for lineages with ancestral functions. Functionally, this would suggest that E2A, while crucial for hematopoietic ancestral functions, was co-opted for lymphoid development and B-cell development.

Until recently, it was hypothesized that the adaptive immune system arose from an evolutionary “big bang” at the speciation of the *Gnathostomata* ^1^. However, advances in genome sequencing of lower *Deuterostomata* has shifted this dogma by describing the presence of adaptive immunity related genes (including RAG, histocompatibility genes and immune type receptors) previously thought to be restricted to the *Gnathostomata* ^21-26,48^. The presence of lymphoid-like cells in lower chordates ^18-20^ further suggests that the lymphoid genetic toolbox was present before the emergence of humoral immunity. Accordingly, the E-protein dependence of the early lymphoid program suggest that the appearance of the full *Gnathostomata* E-protein repertoire was crucial for reinforcing the gene regulatory circuits that drove the emergence and expansion of the hematopoietic lineages constituting humoral immunity.

## Material and Methods

### Animal studies

To generate mice lacking specific E-proteins in the hematopoietic system, Vav-iCre ^57^ was used in combination with conditional (floxed) E2-2 ^58^, HEB ^41^ and E2A ^59^ alleles. Mice were maintained on a C57BL/6 background and analyzed at 8 to 14 weeks of age. Animal studies were approved by the local ethics committee (ethical approval number S16-15).

### Preparation of cells and flow cytometry

Bones, spleen and thymus were dissected, crushed in PBS with 2% FCS and cells were collected after passing through a 70 µm filter. They were then Fc-blocked (CD16/32; 93) and stained with combinations of the antibodies Sca1 (D7), CD105 (MJ7/18), CD41 (MWReg30), CD48 (HM48-1), CD3 (145-2C11), CD4 (RM4-5), CD8 (53-6.7), B220 (RA3-6B2), NK1.1 (PK136), Mac1 (M1/70), Gr1 (RB6-8C5), TER119 (TER-119), CD150 (TCF15-12F12.2), CD117 (2B8, eBioscience), CD127 (A7R34), CD44 (IM7), CD25 (PC61.5, eBioscience), CD19 (1D3, eBioscience), TcRβ (H57-597, eBioscience), TcRγδ (GL3, eBioscience), Ly6C (AL-21), Ly6G (1A8), MHCII (M5/114.15.2), CD11c (N418), PDCA1 (927), Ly6D (49H4), Flt3 (A2F10), IgD (11-26c.2a) and IgM (11/41, eBioscience). All antibodies were purchased from BD Biosciences unless otherwise indicated. For hematopoietic stem and progenitor cell (HSPC) isolation, the cells were subjected to lineage depletion using Dynabeads sheep anti rat IgG (Life Technologies) together with TER119, CD19, CD3, Gr1 and CD11b antibodies. Analysis and cell sorting was performed primarily on an LSR Fortessa and FACSAria IIu (BD Biosciences). Analysis of data was done using the Flowjo 9.9.6 software (Flowjo).

### Phylogenetic analysis

The cDNA and amino acids sequences of the E-proteins from analyzed organisms were obtained through the E-ensembl repository ^60^. See Table S1 for the sequences used in this study. Phylogenetic trees were constructed with MEGA7 ^61^ selecting the Maximum Likelihood method based on the Tamura-Nei model; creating initial tree(s) using the Neighbor-Joining and BioNJ algorithms; and using a Gamma distribution with invariant sites. To assess the support of each node, the tree was bootstrapped 500 times.

### RNA sequencing and analysis

5-10 × 10^3^ cells were FACS-sorted into buffer RLT with β-mercaptoethanol and total RNA extracted using RNeasy Micro Kit (Qiagen, Hilden, Germany) with on-column DNase I treatment. Strand specific RNAseq libraries were prepared using the TotalScript^TM^ RNA-seq kit (Epicentre, Madison, WI) together with custom made Tn5 (transposase). Barcoded libraries were pooled and pair-end sequenced (2×50 cycles) using the Illumina platform (NextSeq500, Illumina, San Diego, CA).

RNAseq reads were mapped using STAR ^62^, reads in exons quantified using HOMER ^63^ and significant changes identified using EdgeR. PCA analysis and display was performed using R (v3.3.3). For details see Supplemental Materials and Methods.

### Pro B cell expansion cultures and fixation

B220^+^ cells were isolated from bone marrow of ER-Cre mice ^64^ using magnetic cell separation (Miltenyi Biotec), expanded for six days in the presence of IL-7 and SCF to obtain pro B-cells. Pro B-cells were subsequently retrovirally transduced with Bcl-2, expanded for seven additional days with 5µM 4-Hydroxytamoxifen in the medium during the last 72 hours. Pro B-cells were fixed using EGS (1.5mM for 30min) in combination with PFA (1% for 10min) and stored as pellets in −80°C.

### ChIP sequencing and analysis

Fixed pro B-cells were thawed, resuspended in SDS lysis buffer, sonicated, Triton X-100 was added and lysates subjected to ChIP by adding dynabeads pre-coupled with either antibodies against E2A, HEB or E2-2. ChIPed chromatin was washed and libraries prepared using reverse crosslinking in conjunction with adapter ligation or via direct amplification of tagmented libraries without prior DNA purification. For details see Supplemental Materials and Methods. Libraries were sequenced using the HiSeq2000 or NextSeq500 platforms (Illumina).

ChIPseq reads were mapped using bowtie2 ^65^. Identification of peaks, peak overlaps and motif enrichment/identification analysis was done using HOMER’s findPeaks, mergePeaks and findMotifsGenome.pl respectively ^63^. Visualization was done using the UCSC genome browser. For details see Supplemental Materials and Methods.

## Data availability

Raw sequencing data are available from the European Nucleotide Archive (ENA) under accession number PRJEB29568 (E-protein knock-out RNAseq and E-protein ChIPseq) and PRJEB20316 (wild-type C57bl/6 RNAseq data).

## Acknowledgements

We thank: Prof. Joakim Dillner and colleagues (Karolinska Institutet) for access to the NextSeq 500 system; Associate Prof. Anders S. Nilsson (Stockholm University) for critical input on the phylogenetic analysis; Karolinska High Throughput Center (KHTC, Karolinska Institutet) for assistance with high-throughput sequencing; the preclinical laboratory at Karolinska University Hospital for access to their facilities; and UPPMAX Next Generation Sequencing Cluster & Storage (UPPNEX) for computational resources. This work was funded by the Swedish Cancer Foundation (Cancerfonden), the Swedish Research Council (VR), the Knut and Alice Wallenberg Foundation (KAW), the Swedish Foundation for Strategic Research (SSF), the European Commission (FP7 PIRG08-GA-2010-276906) and a generous donation by Björn and Lena Ulvaeus.

## Authorship Contributions and conflict of interest statement

TB, CG and RM planned the study; TB, SK, XL, AK, MH, CG, KS and RM performed the experiments; TB, LPP, SK, XL, MH, ADP, CG, KS and RM analyzed the data; CS and DH provided critical insights; CS and RM supervised the research; TB, LPP, CG and RM wrote the manuscript; and all authors reviewed the manuscript before submission.

## Materials & Correspondence

Correspondence and request for material should be addressed to RM.

